# Comparing fMRI inter-subject correlations between groups using permutation tests

**DOI:** 10.1101/370023

**Authors:** Jussi Tohka, Frank E. Pollick, Juha Pajula, Jukka-Pekka Kauppi

## Abstract

Inter-subject correlation (ISC) based analysis is a conceptually simple approach to analyze functional magnetic resonance imaging (fMRI) data acquired under naturalistic stimuli such as a movie. We describe and validate the statistical approaches for comparing ISCs between two groups of subjects implemented in the ISC toolbox, which is an open source software package for ISC-based analysis of fMRI data. The approaches are based on permutation tests. We validated the approaches using five different data sets from the ICBM functional reference battery tasks. First, we created five null datasets (one for each task) by dividing the subjects into two matched groups and assumed that no group difference exists. Second, based on one null dataset, we created datasets with simulated ISC differences of varying size between the two groups. Based on the experiments with these two types of data, we recommend the use of subject-wise permutations, instead of element-wise permutations. The tests based on subject-wise permutations led to correct false positive rates. We observed that the null-distributions should be voxel-specific and not based on pooling all voxels across the brain as is typical in fMRI. This was the case even if studentized permutation tests were used. Additionally, we experimented with an fMRI dataset acquired using a dance movie stimulus for comparison of a group of adult males on the autism spectrum to a matched typically developed group. The experiment confirmed the differences between voxel-based permutation tests and global model based permutation tests.

## 1. Introduction

Inter-subject correlation (ISC) based analysis, originally introduced by Hasson et al. (2004), is a conceptually simple approach to analyze functional magnetic resonance imaging (fMRI) data acquired under naturalistic stimuli such as a movie. In the ISC based analysis of the functional magnetic resonance imaging (fMRI) data, the extent of shared processing across subjects during the experiment is determined by calculating correlation coefficients between the fMRI time series of the subjects in the corresponding brain locations and then averaging the correlation coefficients. ISC maps have been shown to align well with the activation maps of the traditional block design stimuli (Pajula et al., 2012), and it has been applied in a number of studies, e.g., (Hasson et al., 2008; Abrams et al., 2013; Jääskeläinen et al., 2008; Englander et al., 2012; Kauppi et al., 2010; Nummenmaa et al., 2012; Wilson et al., 2007). In addition to localizing shared processing within a group, the ISC methodology has been extended to compare ISCs between two similar, but not identical, stimuli within a single group of subjects (Herbec et al., 2015; Reason et al., 2016), as well as to compare between two groups of subjects experiencing the same stimuli (Hasson et al., 2009; Salmi et al., 2013; Byrge et al., 2015). In this paper, we are interested in the latter scenario. An example application, highlighted also in this paper, is a comparison of the ISCs of participants on the autism spectrum to the ISCs of matched typically developed adults.

A previous study involving the comparison of ISCs between different groups of subjects, Hasson et al. (2009) compared the extent and strength of ISCs of autism and typical groups during free-viewing of a movie. They computed ISC maps separately for both groups as well as between the two groups. To construct the ISC maps, they averaged Fisher z transformed subject-pairwise ISCs within each voxel/region-of-interest. The maps were thresholded based on a maximum value obtained from the identical procedure but using correlations of forward and reversed time courses. Salmi et al. (2013) used a t-statistic to assess difference between the average ISCs of the autistic and typical groups. A permutation test was used to assess statistical significance of the t-statistic. In the test, subjects were randomly exchanged between the groups before re-calculating the t-statistic. A permutation procedure is required because the t-statistic in itself is not distributed according to the t-distribution as the elements of the correlation matrix are not independent. Hence, using a t-test to assess the ISC difference as done in (Byrge et al., 2015), is not appropriate (Chen et al., 2016).

Chen et al. (2016) proposed permutation and bootstrap based methods for statistical hypothesis testing for comparing ISCs between two groups, and Chen et al. (2017) introduced a linear mixed effects model as an alternative to non-parametric ISC hypothesis tests. In particular, Chen et al. (2016) demonstrated that a permutation test exchanging the components of the correlation matrix leads to excessively liberal hypothesis tests while exchanging subjects between the two groups leads to tests with approximately correct false positive rates. In this paper, we will extend the analysis in several ways: 1) we will re-confirm the above mentioned conclusion by new experiments; 2) we will introduce studentized statistics for testing the differences of ISCs between groups; 3) we will show that the voxel-level null-distributions are more appropriate than the image-level (global) null-distributions as even studentized statistics cannot be assumed to be identically distributed across the brain voxels; 4) we will discuss approaches for multiple comparisons correction.

The methods, which we describe, are implemented in the ISC toolbox (Kauppi et al., 2014), which is an open source software package for ISC analysis available at https://www.nitrc.org/projects/isc-toolbox/.

## 2. ISC group comparison

### 2.1 Test-statistic

Let us denote a number of time points in an fMRI time course by *T*, and a number of subjects of *i*th group by *N*_*i*_ (*i* = 1, 2). The fMRI time courses of the voxel *j* from all the subjects in two groups are 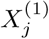 and 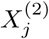, where 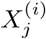 is a *N*_*i*_ *× T* matrix. Further, let 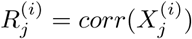 and 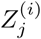 its Fisher’s z-transform ^1^. We denote the element (*n, m*) of 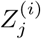 by 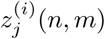, which is the z-transformed correlation coefficient between the time courses of the subjects *m* and *n*. The test statistic for comparing the ISCs between two groups is the difference between the means of z-transformed correlations:

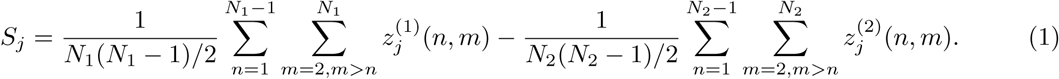

To develop the hypothesis testing for *S*_*j*_, we introduce a matrix 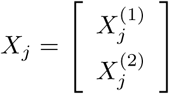 and define *R*_*j*_ = *corr*(*X*_*j*_) and *Z*_*j*_ as the (element-wise) z-transformation of *R*_*j*_. In terms of *Z*_*j*_, we can write *S*_*j*_ in an equivalent
form

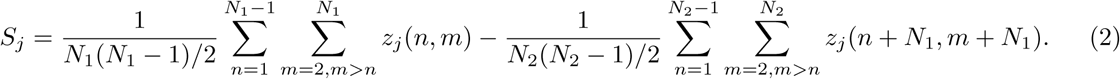

This form will be needed later when developing permutation based inference strategy.

In randomization based hypothesis testing, it is useful to consider studentized test statistics when the interest is in difference between particular parameters, such as means or medians, of two distributions (Chung and Romano, 2013). This is because a permutation test is sensitive to all differences between the two distributions, rather than to the difference in a particular parameter (Chung and Romano, 2013).

This leads us to using studentized test statistics

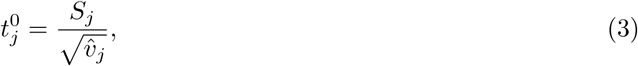

where 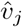 is an estimate of variance of *S*_*j*_. We approximate the variance for each group by

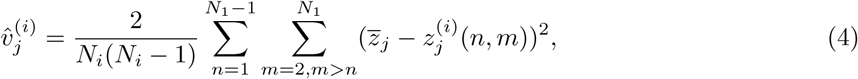

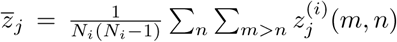 This is a biased variance estimate as it does not account for the dependencies between the elements of the correlation matrix 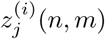. On the other hand, it is an approximation, up to a multiplication by a constant, of the leave-one-subject-out variance defined by Kauppi et al. (2017). We define

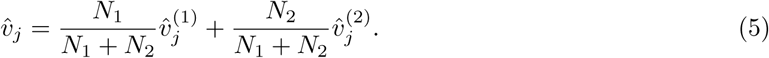

It is important to note that the variance estimates can be also written in terms of the matrix *X*_*j*_ (similarly to the statistic *S*_*j*_ in Eq. (2) above) as this is required for the subject-wise permutation strategy. We further define the test statistic as the SAM statistic (Tusher et al., 2001; Xie et al., 2005)

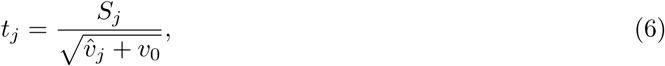

where *v*_0_ is a small positive constant. We will study two versions of this statistic: 1) 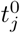, where *v*_0_ = 0 (this is equivalent to Eq. (3)), and 2) 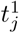, where *v*_0_ is set as the 0.25% percentile of 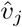 across the brain voxels. The motivation of using this latter statistic is that the variance estimation is regularized, and it is not badly affected by possible outliers. Finally, we emphasize that just normalizing the test statistic values (*S*_*j*_) is not enough for a studentized permutation test, but the variance estimates must be re-computed for each permutation.

### 2.2. Permutation strategies

We study two types of the permutation strategies similarly to Chen et al. (2016):

- **Subject-wise (SW) permutation**. The rows and columns of the matrices *Z*_*j*_ are permuted by a permutation *π* acting on the set 1, 2, …, *N*_1_ + *N*_2_. The row and column indexes are permuted with the same permutation and we denote the permuted matrix as *π*(*Z*_*j*_). Eq. (2), Eq. (3) or Eq. (6) is used to compute the test statistic value after the permutation. This type of permutation strategy necessitates computing also between-group ISCs (i.e., the ISCs of time series of subjects from different groups). This procedure corresponds to random swapping of subjects between the two groups before computing subject-pairwise ISCs. However, it is faster to pre-compute the (z-transformed) correlation matrices and work with them than to compute the test statistics based on time series. A convenient way to denote this permutation is via the permutation matrix *P*_*π*_ corresponding to 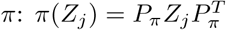.
- **Element-wise (EW) permutation**. The elements of the correlation matrices 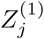 and 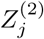 are randomly swapped. In total, there are *N*_1_(*N*_1_-1)/2+*N*_2_(*N*_2_-1)/2 elements in these matrices. After the permutation, the permuted matrices 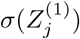 and 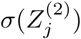 do not have to be proper correlation matrices. In addition, the elements of a correlation matrix are not freely exchangeable and hence the *α*-level of the test based on EW permutations is expected to be overly liberal.

### 2.3. Voxel-null and global-null models

Repeating the permutation procedure *B* times leads to the hypothesis test using one of the above permutation processes. However, an important question remains: should one generate the null-model for each voxel independently (we term these as voxel-null models) or can one assume the same null model for all voxels (we term these as global-null models)? Simplified pseudocodes of these voxel-null and global-null tests are presented in the Algorithms 1 and 2 of Appendix A. In other words, the question is if the test statistics for all voxels are identically distributed (Ge et al., 2003). The global-null approach is widely used in fMRI and, especially, with studentized *t*_*j*_ statistics one might assume that this approach is reasonable. However, as we will show, also *t*_*j*_ statistic values correlate with the average ISC 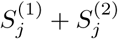, and thus the global-null model suffers from problems. Therefore, we prefer to compute a null model separately for every voxel to obtain a null distribution (and a *p* value) for that voxel. This *p*-value is then Gaussianized using the p-to-Z transform. The disadvantage of this procedure is that it limits the options for multiple comparisons correction (MCC) by essentially ruling out the permutation-based cluster-extent or peak corrections for computational reasons as this would necessitate nested permutation iterations, termed double permutation by Westfall and Young (1993). The Gaussian Random Field (GRF) based corrections (Hayasaka and Nichols, 2003) are still available by turning voxel *p*-values into a *Z*-field by the p-to-Z transform. Similarly, corrections based on the false discovery rate (FDR) are available (Ge et al., 2003). In particular, we implement FDR corrections based on Storey’s procedure (Storey, 2002; Storey and Tibshirani, 2003), which we have found to be better powered than the typical Benjamini-Hochberg procedure (Benjamini and Hochberg, 1995).

### 2.4 Implementation in the ISC toolbox

Our current implementation in the ISC toolbox produces both uncorrected global-null and voxel-null models with the voxel-null model strongly recommended. To form an uncorrected p-value map based on the global-null model, as shown in Algorithm 2, we sample *M* voxels from the brain or gray matter mask, repeat permutations *B* times, and pool the resulting *BM* test statistic values to form the null distribution. The default is to set *B* = 5000 and *M* = 1000.

For MCC, only Storey’s FDR corrections are currently implemented for voxel-null models. However, GRF-based procedures from other software packages can be easily adapted as the ISC toolbox produces Z-map based on voxel-null models. We provide scripts for performing GRF-based corrections based on the FSL function `easythresh` (Flitney and Jenkinson, 2000). For the global-null, we have implemented three MCC strategies 1) FDR-based correction, 2) permutation-based voxel-wise or 3) permutation-based cluster extent correction relying on extreme statistics across the brain (Nichols and Holmes, 2002). To aid the understanding why the voxel-null cannot be directly combined with extreme statistics-based Family-Wise Error (FWE) corrections, we have included pseudo-code of the voxel-wise correction as Algorithm 3 of Appendix A. The cluster extent correction replaces the original voxel-wise statistics by the extents of clusters exceeding a pre-defined cluster defining threshold. As the parametric tests for test-statistics are not available, we set the cluster defining threshold based on raw (i.e., uncorrected) *p*-values approximated by the uncorrected global-null test with a small *B* value (*B* = 250) and including all the brain voxels in each permutation iteration.

The voxel-null models require a large number of permutation iterations, since the MCC cannot be incorporated in the permutation framework and, thus, we have to obtain uncorrected p-values which are accurate especially when they are small. We use the following strategy to speed up the computations: first, we compute *B*_1_ permutation iterations and then fine tune the *p*-values with more iterations for only those voxels that have sufficiently low initial *p*-values (from the first *B*_1_ iterations). This cycle is repeated several times with the *p*-value threshold for fine tuning lowered within every iteration. This procedure saves a substantial amount of computation expense. We recommend to set *B*_1_ to 5000 for the voxel-null models. All the experiments reported in this work are based on the parameter values reported in this subsection.

## 3. Materials and methods

### 3.1. Null experiments

We generated five data sets where we do not expect to see any between group effect in ISC as the groups were based on the matched groups of healthy young subjects. We refer to these sets as null data. For this, we used the fMRI data from 36 healthy young adults (18 men and 18 women; the average age was 28.2 years from the range from 20 to 36 years) during the ICBM functional reference battery (FRB) tasks https://ida.loni.usc.edu/login.jsp?project=ICBM. The ICBM project (Principal Investigator John Mazziotta, M.D., University of California, Los Angeles) is supported by the National Institute of Biomedical Imaging and BioEngineering. ICBM is the result of efforts of coinvestigators from UCLA, Montreal Neurological Institute, University of Texas at San Antonio, and the Institute of Medicine, Juelich/Heinrich Heine University, Germany. We have used the same data earlier in several ISC evaluation experiments (Pajula et al., 2012; Pajula and Tohka, 2014, 2016). The subject numbers, demographics, and the group division can be found in a supplement.

All five FRB tasks were block-design tasks (12 blocks per run (6 off-on blocks) and 3 volumes at the beginning of the run to wait for magnetisation stabilisation), which were highly standardized. In the auditory naming task (AN), subjects were instructed to listen to the description of an object from a sound file and then think their answer silently to the description. In the external order (EO) task, the subjects were presented with four abstract design stimuli followed by a fifth stimulus and required to recall whether the final abstract design was among the four presented previously. In the HA task subjects were instructed to imitate the presented hand configuration with their right hand. In the VG task, the images of certain objects were shown to the subjects on the screen and subjects were instructed to generate a verb associated to the object silently in their mind without saying it aloud. In the OM task, subjects watched an image including a central cross in the middle surrounded by 10 black boxes. Subjects were instructed to concentrate on the central cross and saccade to the surrounding box if it changed white for a moment. After this, they were instructed to return their gaze immediately to the central cross. For a more detailed description of the five tasks, see (Pajula et al., 2012).

The functional data were collected with a 3 Tesla Siemens Allegra fMRI scanner and the anatomical T1 weighted MRI data with an 1.5 Tesla Siemens Sonata scanner. The TR/TE times for the functional data were 4 s/32 ms, flip angle 90 degree, pixel spacing 2 mm and slice thickness 2 mm. The parameters for the anatomical T1 data were 1.1 s/4.38 ms, 15 degree, 1 mm and 1 mm, correspondingly. The fMRI data were preprocessed (including motion correction, stereotactic registration, temporal high-pass filtering with a cutoff period of 60s, spatial filtering with 5mm isotropic kernel) using FSL as described in (Pajula et al., 2012).

We divided 36 subjects randomly into two groups of 18 subjects so that the groups were age and sex matched. In this setting, with highly similar groups, there should be no difference between the two groups. We further verified that no group differences existed by using the standard general linear model based hypothesis test as implemented in FSL’s (FMRIB’s Software Library, www.fmrib.ox.ac.uk/fsl) FEAT (FMRI Expert Analysis Tool) Version 6.00. Higher-level analysis was carried out using FLAME (FMRIB’s Local Analysis of Mixed Effects) stage 1 and stage 2 (Beckmann et al., 2003; Woolrich et al., 2004; Woolrich, 2008). The standard analysis is different from the ISC-based analysis in that it requires reference time courses whereas the ISC-based analysis does not. The false positive rates for this analysis are presented in the Appendix B.

### 3.2. Simulated ISC differences

Starting from the null data for the AN task, described in the previous sub-section, we generated data with simulated ISC differences between the two groups *G*_1_ and *G*_2_. Let 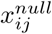 be de-meaned and normalized (to the unit variance) the time series of the subject *i* at the voxel *j* of the null data. Then, the simulated data was defined as

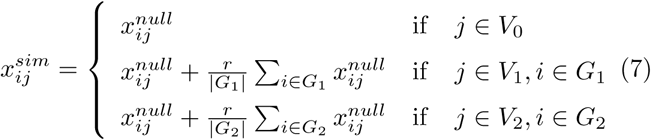

where *r >* 0 is a parameter related to the size of the effect, *V*_0_ is a set of voxels where the two groups are assumed to have no differences, *V*_1_ is s set of voxels where ISC of *G*_1_ is assumed to be greater than ISC of *G*_2_, and *V*_2_ is s set of voxels where ISC of *G*_2_ is assumed to be greater than ISC of *G*_1_. We simulated the data with three different values for *r*: 0.1, 0.25, and 0.5. *V*_1_ was to selected to be the union of regions 29 (Cingulate, anterior division) and 42 (Central opercular cortex) of the Harvard-Oxford atlas. *V*_2_ was to selected to be the region 22 (lateral occipital cortex) of the Harvard-Oxford atlas. *V*_1_ contained 2490 voxels, *V*_2_ 5134 voxels, and *V*_0_ 206376 voxels. We note that we did not to try to model any task with this simulation. The groups were defined as in Section 3.1.

### 3.3. Autism spectrum experiment

Additionally, we have applied the method to compare ISC of 8 male adults on the autism spectrum (ASD) to 8 age and IQ matched typically developed (TD) male adults while they viewed a 90 second clip of a solo ballet dance. The TD group was comprised of 8 individuals with an average age of 27.5 (SD 7.4), and the ASD group included 8 individuals with an average age of 28.5 (SD 8.1). The TD group had an average Autism Quotient (AQ) score of 12.3 (SD 5.5) (N=7) and an average Intelligence Quotient (IQ) score of 119.0 (SD 7.7). The ASD group had an average AQ of 38.9 (SD 7.1) (N=7) and an average IQ score of 118.9 (SD 6.0). All were right handed as assessed by the Edinburgh handedness inventory. Participants were recruited from the participant database at the School of Psychology, University of Glasgow. None of the participants had experience in practicing ballet dance and none regularly watched dance performances. Ethical permission for the study was obtained from the Greater Glasgow and Clyde National Health Service ethics board.

While in the scanner, all participants viewed three dance videos and in the present analysis we examine just one. This stimulus was a video (60 fps, 1280 by 720 resolution) of a Romantic ballet dance (Giselles solo dance in Act II of Giselle), 90 seconds in duration. The video was also converted to black and white, the ballerinas face was blurred out and there was no associated audio track. Stimulus presentation was controlled by Presentation software (Neurobehavioural systems, Inc). Before beginning the experiment, participants were instructed to simply relax and enjoy watching the dances while being scanned.

Data were acquired from a single functional T2*-weighted acquisition (EPI, TR 2000 ms; TE 30 ms; 32 Slices; 3mm^3^ voxels; FOV of 210, imaging matrix of 70 ×70) using a 3T Tim Trio Siemens scanner. The run took 270 seconds with a total of 90 seconds for each dance presentation. There were 8 seconds of blank at the beginning and 36 seconds at the end of the run and 16 seconds of blank between the first and second as well as the second and third dance presentation. The Romantic style dance chosen for analysis occurred randomly in either the first or second position. An anatomical scan was performed at the end of the scanning session that comprised a high-resolution T1-weighted anatomical scan using a 3D magnetization prepared rapid acquisition gradient recalled echo (ADNI-MPRAGE) T1-weighted sequence (192 slices; 1mm cube isovoxel; Sagittal Slice; TR = 1900 ms; TE = 2.52; 256 × 256 image resolution). The fMRI data were preprocessed in Brain Voyager QX (Vers.2.6, Brain Innovation B.V., Maastricht, Netherlands). This included: 3D Motion Correction with Trilinear/sinc interpolation, slice scan-time correction, linear removal, and high-pass filtering with cutoff set to 1 cycle. Spatial smoothing with a Gaussian kernel of 6 mm FWHM was also applied. This was followed by normalization of functional scans into common Talairach space, and co-registration of functional and anatomical data. Finally, the functional data were trimmed using Matlab to obtain the 45 volumes (90 seconds) for each dance, used later for ISC analysis.

## 4. Results

### 4.1. Null experiment

The results of the null experiment are listed in Tables 1 and 2. In the Tables, the fractions of significant voxels (uncorrected) across the gray matter mask are displayed when the *α*-level is varied. We call this fraction as (observed) false positive rate (FPR). It should be as close as possible to the nominal *α*-level (0.05, 0.01, or 0.001 in the tables). As the tables show, the subject-wise permutations led to the approximately correct observed FPRs. However, the element-wise permutations produced too liberal p-values, i.e., the observed FPRs were much higher than the nominal *α*-level. These results agree with Chen et al. (2016). The tables also list the correlations between the Z-transformed p-value for the group difference and the average ISC of the two groups. This correlation should be as close to zero as possible as the average ISC between two groups should be independent of the magnitude of ISC difference when no difference is expected. As the tables show, only the subject-wise voxel-null models displayed desirable behavior whereas subject-wise global-null models displayed correlations from 0.33 (with AN task) to 0.57 (with HA task). The studentization of the permutation tests reduced the correlation but did not eliminate it. The high correlation values indicated that voxels with high average ISC were much more prone to display a significant ISC difference by chance than the voxels with low average ISC. The undesirable performance of the global-null models is visualized in Figure 1. As can be seen in the Figure, the global-null model (thresholded at *p* < 0.05, SW-permutations with studentization) produced large clusters of significant voxels. This performance was not expected for a null experiment and could be problematic especially when cluster extents are studied for MCC. Instead, the voxel-null model produced a comparison map which appeared more realistic with small clusters of significant voxels across the brain (see Fig. 1).

**Table 1:**
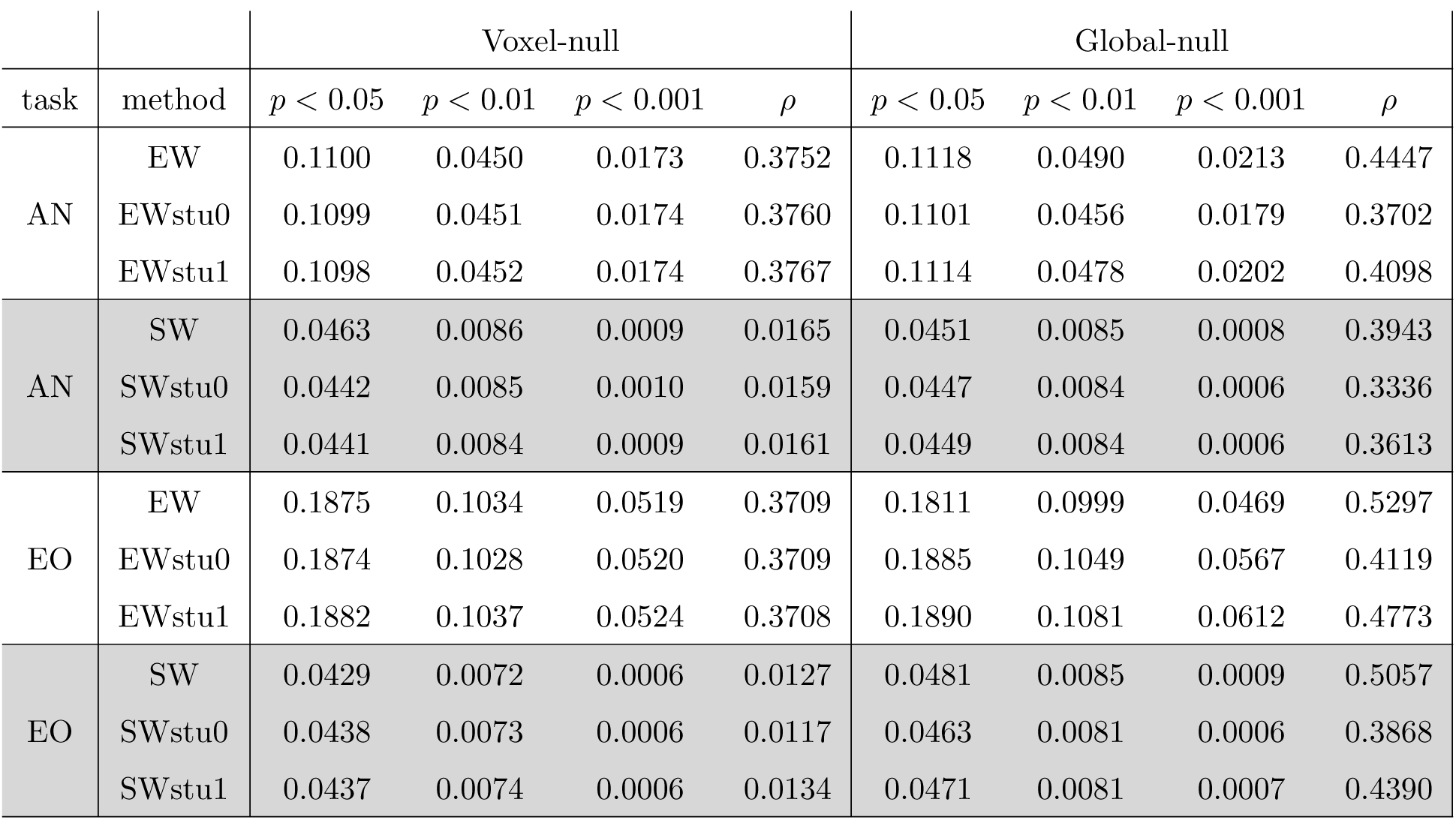
The results of the null experiment with the AN and EO stimuli of the ICBM-FRB. The columns *p* < 0.05, *p* < 0.01, and *p* < 0.001 list the fraction of the significant voxels at each *α* level (false positive rate). The closer the value is to the nominal *α*-level, the better the result is. The columns *ρ* list the correlation between the Z-transformed p-value for the group difference and the average ISC of the two groups. Lower the absolute correlation, the better the result. EW and SW are the permutation strategies, stu0 is the studentized statistic with *v*_0_ = 0, stu1 is the studentized statistic with *v*_0_ as 0.25 % quantile of voxel-wise variances across the mask (see Section 2.1.). Only SW permutations with voxel-wise null models produced satisfactory results, independent of the studentization strategy.

**Table 2:**
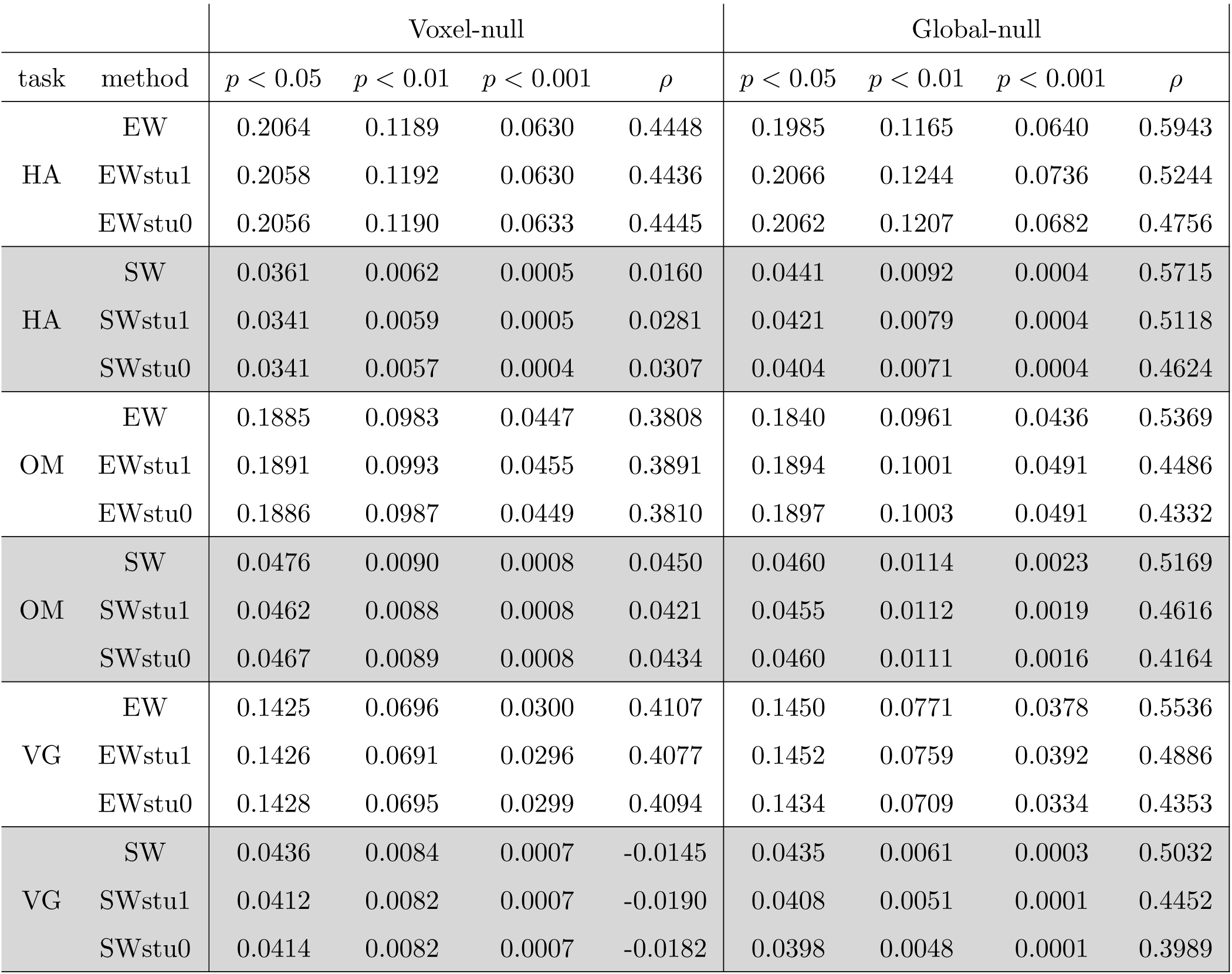
The results of the null experiment with the HA, OM, and VG stimuli of the ICBM-FRB. The columns *p* < 0.05, *p* < 0.01, and *p* < 0.001 list the fraction of the significant voxels (false positive rate) at each *α* level. The closer the value is to the *α*-level, the better the result is. The columns *ρ* list between the Z-transformed p-value for the group difference and the average ISC of the two groups. Lower the absolute correlation, the better the result. EW and SW are the permutation strategies, stu0 is the studentized statistic with *v*_0_ = 0, stu1 is the studentized statistic with *v*_0_ as the 0.25% quantile of voxel-wise variances across the mask (see Section 2.1). Only SW permutations with voxel-wise null models produced satisfactory results, independent of the studentization strategy.

| task | method | Voxel-null |  |  |  | Global-null |  |  |  |
| --- | --- | --- | --- | --- | --- | --- | --- | --- | --- |
| | | $p < 0.05$ | $p < 0.01$ | $p < 0.001$ | $\rho$ | $p < 0.05$ | $p < 0.01$ | $p < 0.001$ | $\rho$ |
| HA | EW | 0.2064 | 0.1189 | 0.0630 | 0.4448 | 0.1985 | 0.1165 | 0.0640 | 0.5943 |
|  | EWstu1 | 0.2058 | 0.1192 | 0.0630 | 0.4436 | 0.2066 | 0.1244 | 0.0736 | 0.5244 |
|  | EWstu0 | 0.2056 | 0.1190 | 0.0633 | 0.4445 | 0.2062 | 0.1207 | 0.0682 | 0.4756 |
| HA | SW | 0.0361 | 0.0062 | 0.0005 | 0.0160 | 0.0441 | 0.0092 | 0.0004 | 0.5715 |
|  | SWstu1 | 0.0341 | 0.0059 | 0.0005 | 0.0281 | 0.0421 | 0.0079 | 0.0004 | 0.5118 |
|  | SWstu0 | 0.0341 | 0.0057 | 0.0004 | 0.0307 | 0.0404 | 0.0071 | 0.0004 | 0.4624 |
| OM | EW | 0.1885 | 0.0983 | 0.0447 | 0.3808 | 0.1840 | 0.0961 | 0.0436 | 0.5369 |
|  | EWstu1 | 0.1891 | 0.0993 | 0.0455 | 0.3891 | 0.1894 | 0.1001 | 0.0491 | 0.4486 |
|  | EWstu0 | 0.1886 | 0.0987 | 0.0449 | 0.3810 | 0.1897 | 0.1003 | 0.0491 | 0.4332 |
| OM | SW | 0.0476 | 0.0090 | 0.0008 | 0.0450 | 0.0460 | 0.0114 | 0.0023 | 0.5169 |
|  | SWstu1 | 0.0462 | 0.0088 | 0.0008 | 0.0421 | 0.0455 | 0.0112 | 0.0019 | 0.4616 |
|  | SWstu0 | 0.0467 | 0.0089 | 0.0008 | 0.0434 | 0.0460 | 0.0111 | 0.0016 | 0.4164 |
| VG | EW | 0.1425 | 0.0696 | 0.0300 | 0.4107 | 0.1450 | 0.0771 | 0.0378 | 0.5536 |
|  | EWstu1 | 0.1426 | 0.0691 | 0.0296 | 0.4077 | 0.1452 | 0.0759 | 0.0392 | 0.4886 |
|  | EWstu0 | 0.1428 | 0.0695 | 0.0299 | 0.4094 | 0.1434 | 0.0709 | 0.0334 | 0.4353 |
| VG | SW | 0.0436 | 0.0084 | 0.0007 | -0.0145 | 0.0435 | 0.0061 | 0.0003 | 0.5032 |
|  | SWstu1 | 0.0412 | 0.0082 | 0.0007 | -0.0190 | 0.0408 | 0.0051 | 0.0001 | 0.4452 |
|  | SWstu0 | 0.0414 | 0.0082 | 0.0007 | -0.0182 | 0.0398 | 0.0048 | 0.0001 | 0.3989 |

**Figure 1:**
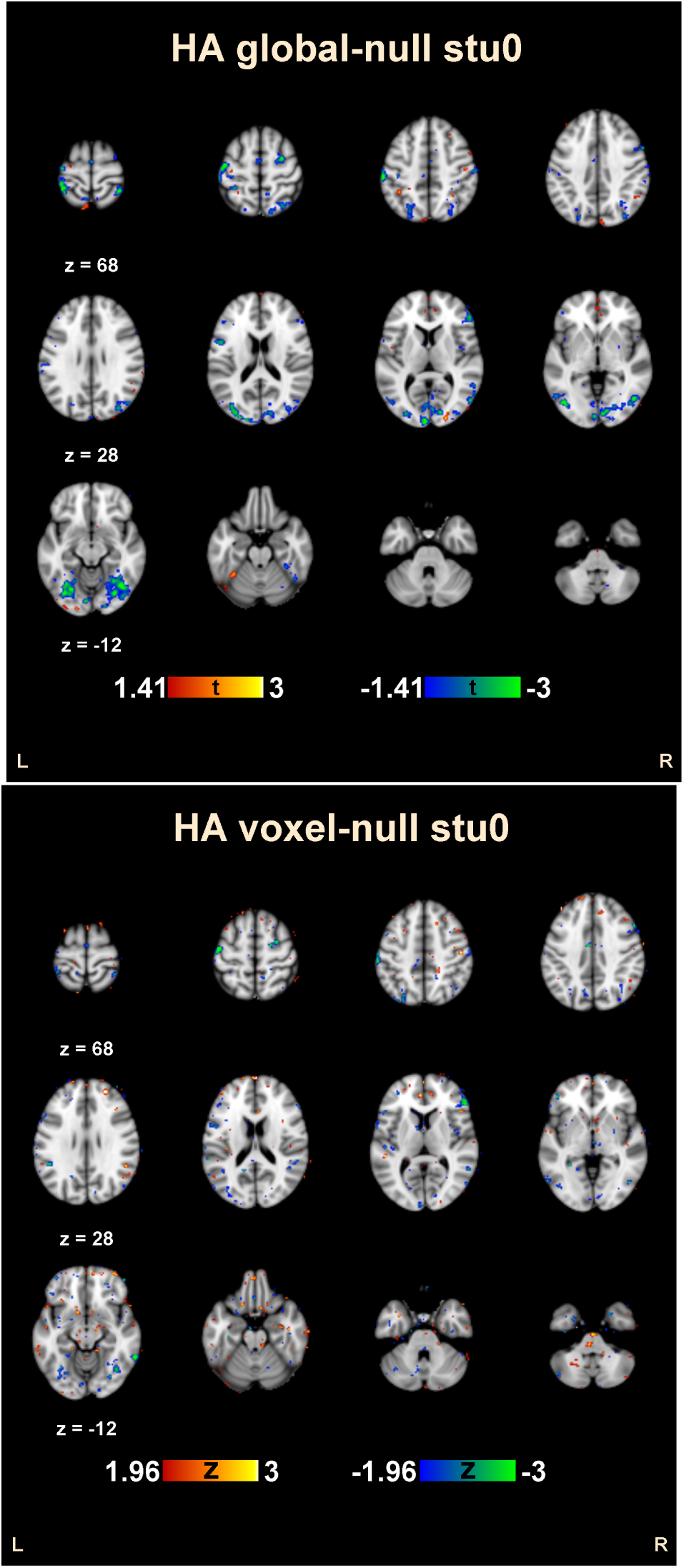
Uncorrected statistics maps of the HA task in the null experiment thresholded at *p* < 0.05 for global-null (upper panel) and voxel-null (lower panel) models. The subject-wise permutations with studentization was applied. The global-null model produced large clusters of significant voxels unlike the voxel-null model, whose statistic map appears more realistic for a null-experiment where differences between groups are expected to be by chance.

### 4.2. Simulated ISC differences

The true positive rates (TPRs) of the simulated ISC experiment are listed in Tables 3. We have only listed the results of the SW permutations that succeeded in maintaining the nominal *α*-level. For these tests, the observed FPRs were well in line with nominal rates also in this experiment and they are not displayed. We have also included the TPR resulting from thresholding of the voxel-wise statistics at the FDR corrected level of 0.05. FDR correction was implemented using Storey’s procedure (Storey and Tibshirani, 2003). Table 4 lists the minimum q-values from the FDR adjustment as well as area under the receiver operating characteristic curve (AUC) and area under the precision recall curve (AUCPR) (Boyd et al., 2013). These quantities are useful for evaluation of binary classification allowing the performance assessment at a range of *p*-value thresholds instead of fixing a single threshold. (The classes are: 1) the voxels that are different between the groups and 2) the voxels where no difference should exist). AUCPR is better of the two in cases where the classes are imbalanced, i.e., contain very different number of voxels, where AUC can fail to depict the performance difference of different methods properly (Boyd et al., 2013).

**Table 3:** The results of the simulated ISC experiment. The columns *p* < 0.05, *p* < 0.01, and *p* < 0.001 list the true positive rate (TPR, the fraction of voxels detected as different between the groups of the truly different voxels between the groups) at each *α* level. The column *q* < 0.05 lists the TPR when the threshold was 0.05 FDR corrected. *r* is the *r*-parameter used in simulation of ISC differences (Eq. (7)). Other notation is as in Table 1.

| r | method | Voxel-null |  |  |  | Global-null |  |  |  |
| --- | --- | --- | --- | --- | --- | --- | --- | --- | --- |
| | | $p < 0.05$ | $p < 0.01$ | $p < 0.001$ | $q < 0.05$ | $p < 0.05$ | $p < 0.01$ | $p < 0.001$ | $q < 0.05$ |
| 0.10 | SWstu1 | 0.1060 | 0.0298 | 0.0041 | 0 | 0.1086 | 0.0138 | 0.0001 | 0 |
| 0.10 | SWstu0 | 0.1051 | 0.0292 | 0.0046 | 0 | 0.1091 | 0.0177 | 0.0004 | 0 |
| 0.10 | SW | 0.1103 | 0.0306 | 0.0042 | 0 | 0.1038 | 0.0106 | 0 | 0 |
| 0.25 | SWstu1 | 0.4124 | 0.2074 | 0.0637 | 0 | 0.4534 | 0.1302 | 0.0063 | 0 |
| 0.25 | SWstu0 | 0.4154 | 0.2067 | 0.0627 | 0 | 0.4587 | 0.1579 | 0.0122 | 0 |
| 0.25 | SW | 0.4244 | 0.2116 | 0.0640 | 0 | 0.4461 | 0.0961 | 0.0024 | 0 |
| 0.50 | SWstu1 | 0.9003 | 0.7626 | 0.5129 | 0.5077 | 0.9259 | 0.7079 | 0.1638 | 0 |
| 0.50 | SWstu0 | 0.9011 | 0.7632 | 0.5135 | 0.5092 | 0.9279 | 0.7467 | 0.2594 | 0.0092 |
| 0.50 | SW | 0.9032 | 0.7631 | 0.5127 | 0.5092 | 0.9234 | 0.6400 | 0.0742 | 0 |

**Table 4:**
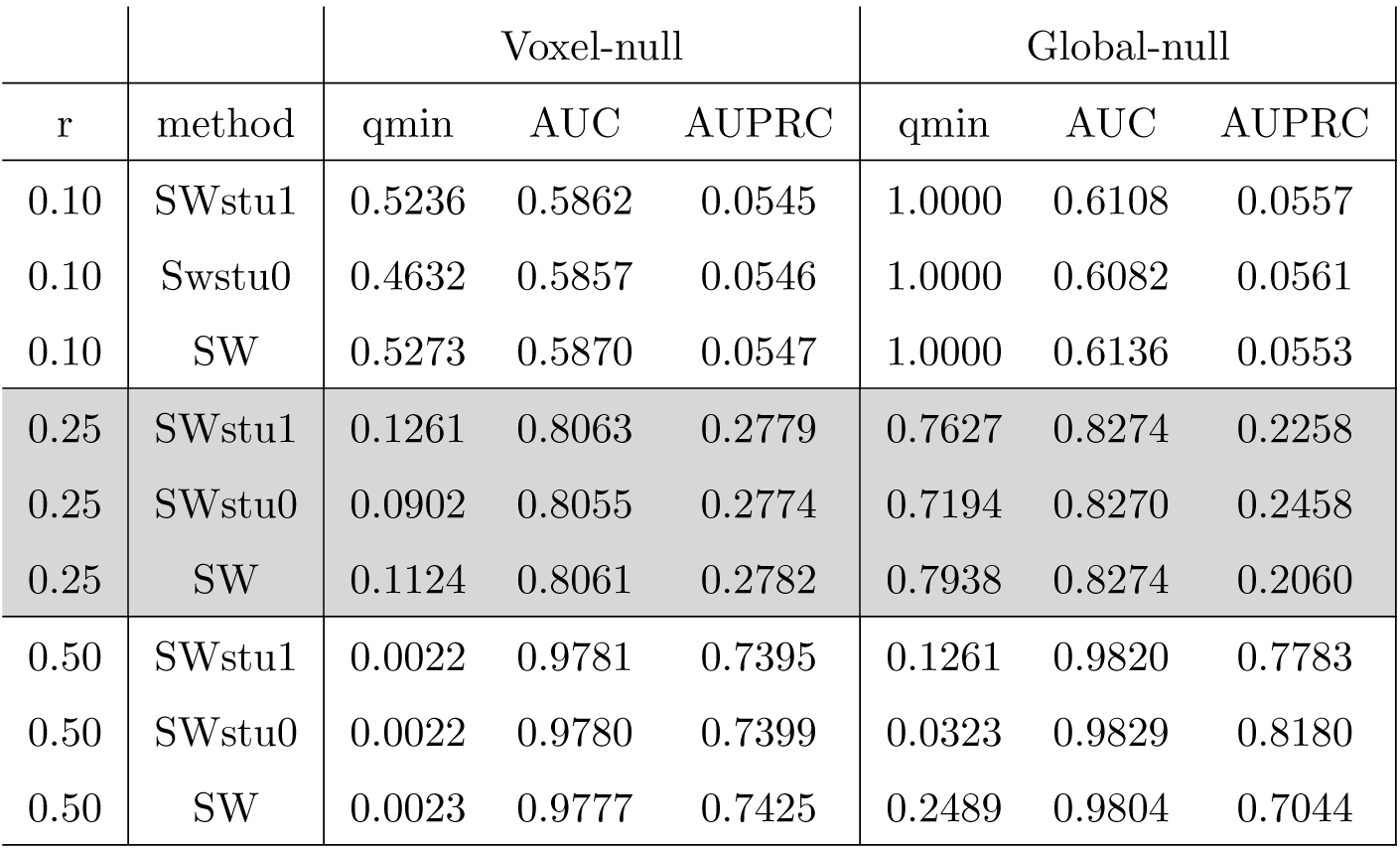
The results of the simulated ISC experiment. The column qmin is the smallest q-value within the gray matter mask. AUC (AUCPR) is the AUC (AUCPR) between voxels with simulated group difference and the voxels which should be similar between the groups. *r* is the *r*-parameter used in simulation of ISC differences (Eq. (7)). Other notation is as in Table 1.

Comparison of TPRs between voxel-null and global-null models in Table 3 reveals that global-null models have difficulties in attaining small p-values. At the threshold *p* < 0.05, the global-null appeared as more powerful than voxel-null, but at lower (and more interesting) p-thresholds this relation was overturned, very markedly in the case of *p* < 0.001. We hypothesize that this phenomenon follows from the dependence of global-null p-values on underlying mean ISC values, which was demonstrated with the null-experiments in Section 4.1. This is also visible in Fig. 2, where the results of voxel-null and global-null models are compared to the ground-truth group differences at different *α*-thresholds.

**Figure 2:**
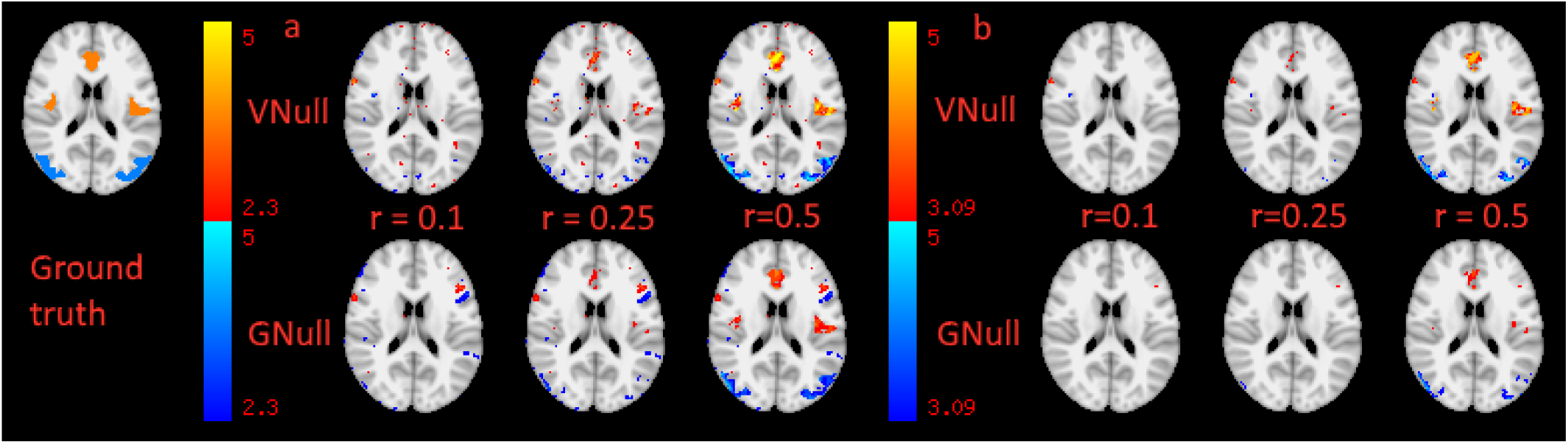
The detected group differences of voxel-null (VNull) and global-null (GNull) models compared to the ground-truth group differences when the threshold *p* = 0.01 (panel a) when the threshold *p* = 0.001 (panel b). Subject-wise permutations with studentization was used. The values shown are z-transformed p-values for both global-null and voxel-null models. An axial slice at MNI coordinate *z* = 18*mm* is shown. Especially, when the threshold is *p* = 0.001 voxel-null model was better powered.

Interestingly, in terms of AUC, the global-null appeared better than the voxel-null. However, in terms of AUCPR, the superiority of the models depended on the value of the power parameter *r* used in the data generation (see Section 3.2) as shown in Table 4. From Tables 3 and 4, it can be observed that the studentization did not markedly alter the TPRs, AUCs, or AUCPRs in the case of voxel-null based models while with the global-null models, the studentization was clearly useful and helped to correct some of the disadvantages of the global-null models. Moreover, studentization without regularization was preferable.

We observed (Table 3) that correcting with FDR threshold of 0.05 only revealed differences when the power parameter had its highest value. The minimal *q*-values in Table 4 reveal that the studentization without regularization might be useful also with voxel-null models as the smallest *q*-values were the smallest.

### 4.3. Multiple comparison correction options

With the simulated differences dataset of Section 3.2, we assessed the following MCC strategies: 1) cluster-extent based correction using a permutation test operating on assumption of the global-null model (gloCE-perm), 2) cluster-extent based correction of z-transformed voxel-null p-values using GRFs as implemented in FSL’s `easythresh`-function (Flitney and Jenkinson, 2000) (voxCE-GRF) and 3) correction based Storey’s FDR procedure acting on voxel-null p-values (voxFDR). For 1), the cluster defining threshold of *p* = 0.01 was considered. For voxCE-GRF we considered cluster definition thresholds of *p* = 0.01 and *p* = 0.001. The first one corresponds (approximately) to the default value used by FSL, but several studies recommend a more conservative primary threshold (Woo et al., 2014). The raw p values were Z-transformed before applying FSL’s `easythresh`. A cluster extent with *p* < 0.05 was considered significant. With FDR correction, a voxel with *q* < 0.05 was considered to be significant.

The true ISC differences in the dataset of Section 3.2 appear in five continuous clusters (Cingulate, anterior division, left and right Central opercular cortex, and left and right Lateral occipital cortex). We measured how many of these five clusters a method could detect and how many false positive clusters a method would detect. We considered a cluster-extent based method to correctly identify a cluster of ISC differences if the overlap between the detected supra-threshold cluster and the true cluster was larger than 50 %. FDR was deemed to correctly identify a cluster if it detected a single voxel within the true cluster. A significant cluster with no overlap with any of the five true clusters, was deemed as false positive. With FDR correction, we instead chose to compute the fraction false positive voxels among positive voxels, i.e., the observed FDR, which should be close to the nominal FDR-level.

The cluster detection rates are listed in Tables 5. Only SW permutations were assessed. The GRF-based cluster extent correction with the primary threshold of *p* = 0.01 identified one false positive cluster when *r* equaled 0.1. Otherwise, no false positive clusters were detected. The observed FDR was 0.044 when *r* = 0.5, otherwise no voxels passed the *q* < 0.05 threshold. The results were equal for all studentization options. As shown in Table 5, the GRF-based cluster extent correction (voxCE-GRF) was the most powerful option in detecting clusters of ISC difference. The more conservative cluster definition threshold (*p* = 0.001) is recommended as it was slightly less powerful but identified no false positive clusters. FDR correction performed well when the simulated ISC difference was largest, but failed to reveal any differences for smaller ISC differences. The permutation-based cluster extent correction identified only the two largest clusters (left and right Lateral occipital cortex) when the ISC difference was the largest, i.e., when *r* = 0.5.

**Table 5:** The number of true clusters detected for multiple comparison options. 5 is the best result indicating all true clusters detected. See the text for method abbreviations. After CE-GRF acronym, the cluster definition threshold is given. Only subject-wise permutations were assessed and all studentization options gave the same results. CE-GRF-0.01 produced one false positive cluster when *r* was 0.25. Other cluster extent based methods produced no false positive clusters. FDR detected no differences for *r* = 0.1, *r* = 0.25. When *r* = 0.5, the observed FDR matched well with the nominal FDR of 0.05.

| r | gloCE-perm | voxCE-GRF 0.01 | voxCE-GRF 0.001 | voxFDR |
| --- | --- | --- | --- | --- |
| 0.1 | 0 | 1 | 1 | 0 |
| 0.25 | 0 | 4 | 3 | 0 |
| 0.5 | 2 | 5 | 5 | 5 |

### 4.4 Autism spectrum experiment

The results of the comparison maps (both voxel null-model and global-null model, studentized with *v*_0_ is set as the 0.25% percentile of 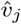 across the brain voxels) thresholded at *p* = 0.01 (uncorrected) showed several brain regions that differed between the groups. The voxel-null results are shown in Fig. 3, and ignoring small clusters less than 108 *mm*^3^, revealed 6 regions where the ISC map was greater for the ASD group, and 12 regions where the ISC map was greater for the TD group. The global-null results are shown in Fig. 4, and ignoring small clusters less than 108 *mm*^3^, revealed 3 clusters where there the ISC map was greater for the ASD group, and 11 regions where the ISC map was greater for the TD group. These results, showing a greater number of regions where ISC is greater for the TD group than the ASD group, are consistent with previous reports of less ISC in autism (Hasson et al., 2009; Salmi et al., 2013). Overlapping clusters for the voxel- and global-null results were found for the contrast *ASD > TD* in the left middle frontal gyrus and for the contrast *TD > ASD* in right posterior cingulate, right precuneus, right middle frontal gyrus, right precentral gyrus and left parahippocampal gyrus. Both the voxel-null and global-null model results were submitted to cluster thresholding using FSL’s easyhtresh with a cluster determining threshold of p=0.001 (Flitney and Jenkinson, 2000). The only cluster to survive was from the voxel-null model for the contrast *TD > ASD* and revealed a cluster of 567 mm^3^ at right posterior cingulate (peak at Talairach coordinates 4, −42, 25). The posterior cingulate can be considered a hub region, with dense connections to other brain regions (Leech and Sharp, 2013; van den Heuvel and Sporns, 2013) and the involvement of hub areas in watching dance has been discussed in previous examinations of watching dance using ISC (Pollick et al., 2018). One potential basis for the lower ISC in posterior cingulate in the autism group is that the posterior cingulate has been shown to have idiosyncratic resting state functional connectivity in autism (Nunes et al., 2018). Finally, as the null-experiment results (see Section 4.1) support the use of the voxel-null model instead of the global-null model, we recommend using the voxel-based null model if there is no other reason to prefer the global-null instead.

**Figure 3:**
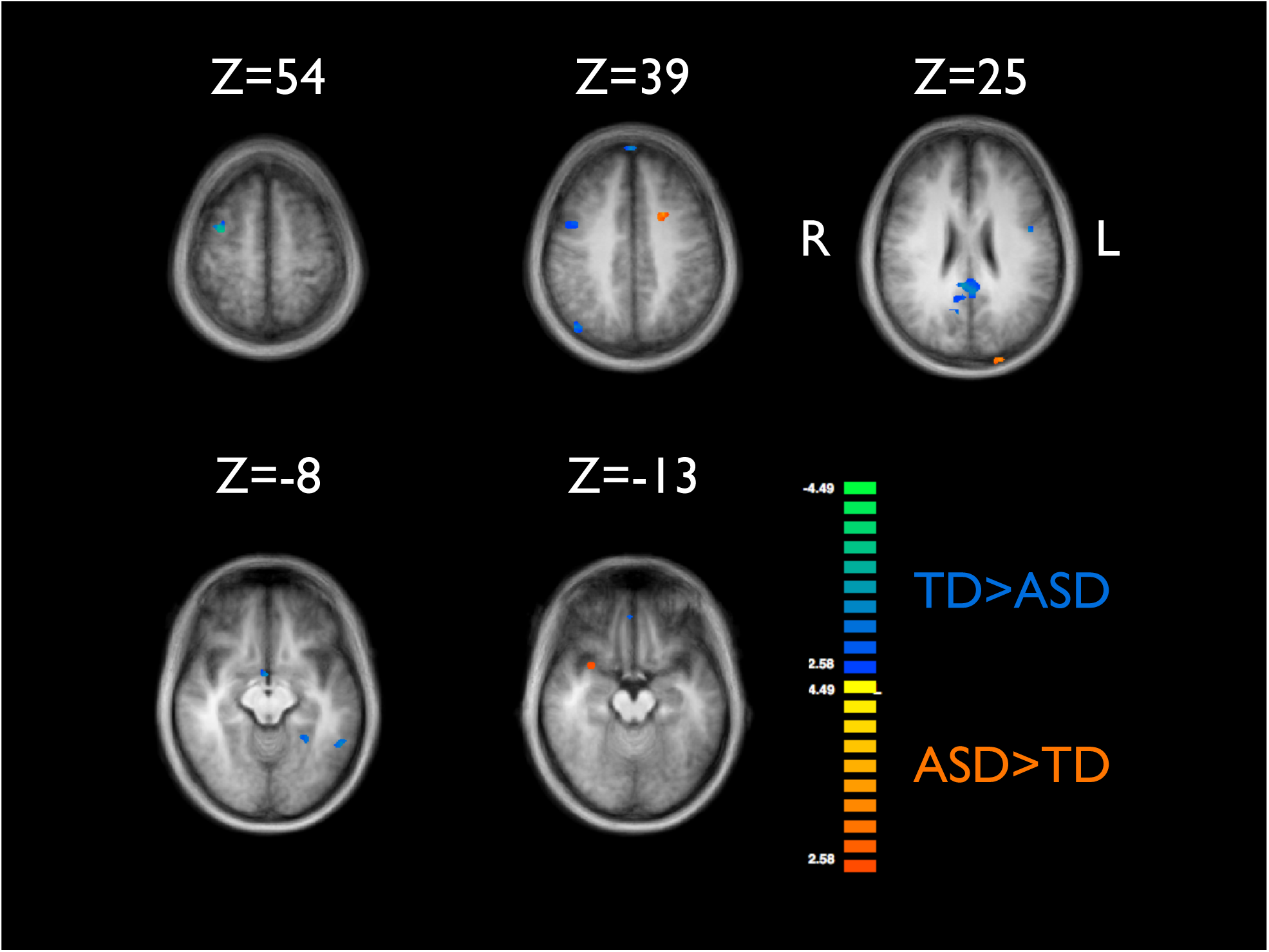
Uncorrected comparison map from the autism spectrum experiment, thresholded at *p* = 0.01. Voxel-null model, studentized with *v*_0_ is set as the 0.25% percentile of 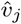 across the brain voxels.

**Figure 4:**
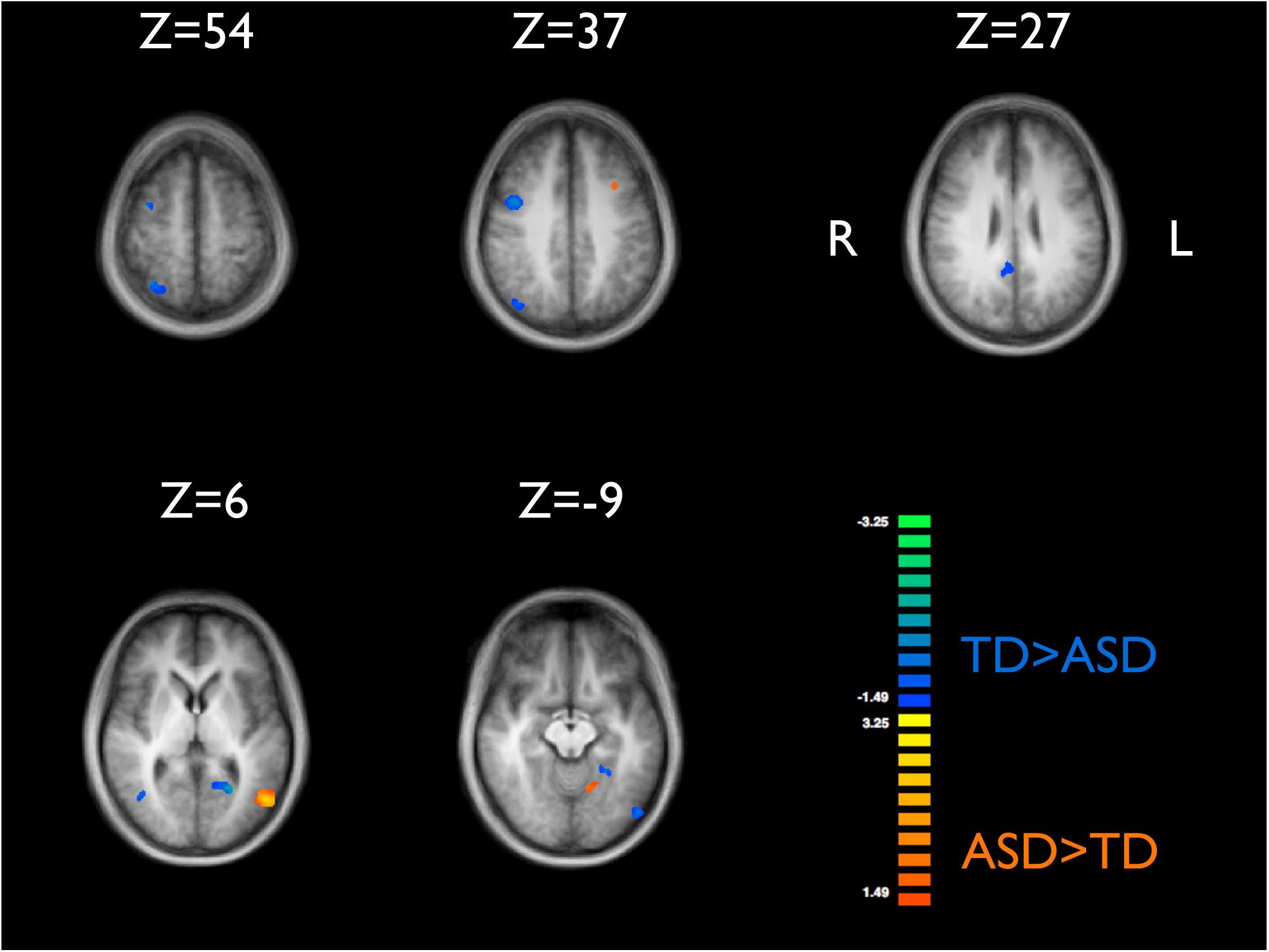
Uncorrected comparison map from the autism spectrum experiment, thresholded at *p* = 0.01. Global-null model, studentized with *v*_0_ is set as the 0.25% percentile of 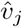 across the brain voxels.

## 5. Discussion

We have presented and studied the ISC group comparison approaches using permutation tests. We have verified that the permutation-based ISC group comparison results in approximately correct type I error rates. Our results agreed with Chen et al. (2016) in that the element-wise permutations led to an unacceptably high false positive rate. We thus strongly recommend a subject-wise permutation strategy. We have presented results that discourage the use of global (across-brain) null-models, and instead we recommend to generate null-models independently for each voxel. The studentization of the test statistic was found to improve the performance of the global-null-models but not to cure their deficiencies entirely. Further, we demonstrated that the GRF-based cluster extent correction when coupled with voxel-null models is a viable alternative for multiple comparison correction.

We have based our conclusions on the experiments that were done on null data sets, where we expected not to find any group difference as the groups were matched, as well as experiments on data sets that contained a simulated ISC difference between the two groups. While Chen et al. (2016) demonstrated the importance of subject-wise rather than element-wise permutations in ISC group comparisons, they based their analysis on simple, purely synthetic data. However, in ISC analysis, this kind of synthetic data fails to reproduce some of the main characteristics of ISCs seen in actual fMRI experiments such as the increase of variance of subject-pair-wise ISCs with the increase of mean ISCs (Kauppi et al., 2017). Thus, we argue that basing the analysis on advanced simulations built on top of actual fMRI data is a more appropriate strategy than using purely synthetic data.

An essential and perhaps surprising finding of this study was that the use of global (i.e., same for every voxel) null-models often led to poor results as the hypothesis of identical test statistic distributions across the voxels does not hold. This finding held even if studentized permutation tests were applied. Thus, using voxel-specific null models seemed to have definite advantages over more typical global-null models. Although not typical, voxel specific null models are not new in neuroimaging. They are used, for example, in the FSL’s FLAME1+2 mixed-effect analysis (Woolrich et al., 2004). Also, in microarray data analysis, gene-specific (corresponding to our voxel-null models) vs. global (all genes together) null models have been contrasted (Ge et al., 2003; Dudoit et al., 2003).

The limitation of voxel-specific models is that the permutation based options for multiple comparisons correction are no longer available without a considerable increase of the computation time as they involve two nested rounds of resampling (see Westfall and Young (1993); Ge et al. (2003)). Of note is that even the faster version of the double permutation method of Ge et al. (2003) would lead to a prohibitive increase of the computation time in the ISC analysis. Regarding multiple comparison correction options, this study leads to recommend Gaussian Random Field based cluster extent correction on top of Z-maps generated based on voxel-null models. However, in simulations the activated areas were continuous clusters with large extents, so that the simulations may have been too well-suited for GRF based cluster extent correction. Thus, in some cases, voxel-level FDR may be a better criterion for multiple comparisons correction. Also, we repeat the warning voiced by many others (e.g., Eklund et al. (2016); Woo et al. (2014)) that GRF-based cluster extent correction should be used with a conservative primary threshold and the widely used value corresponding to *p* = 0.01 may be too lenient. As we demonstrated, permutation-based cluster extent correction performed poorly due to its reliance on global null-models.

The methods presented in this study are implemented in the ISCtoolbox software package https://www.nitrc.org/projects/isc-toolbox/, which is freely available under an open-source licence.

### Algorithm 1 Voxel-null test with SW permutations (uncorrected)

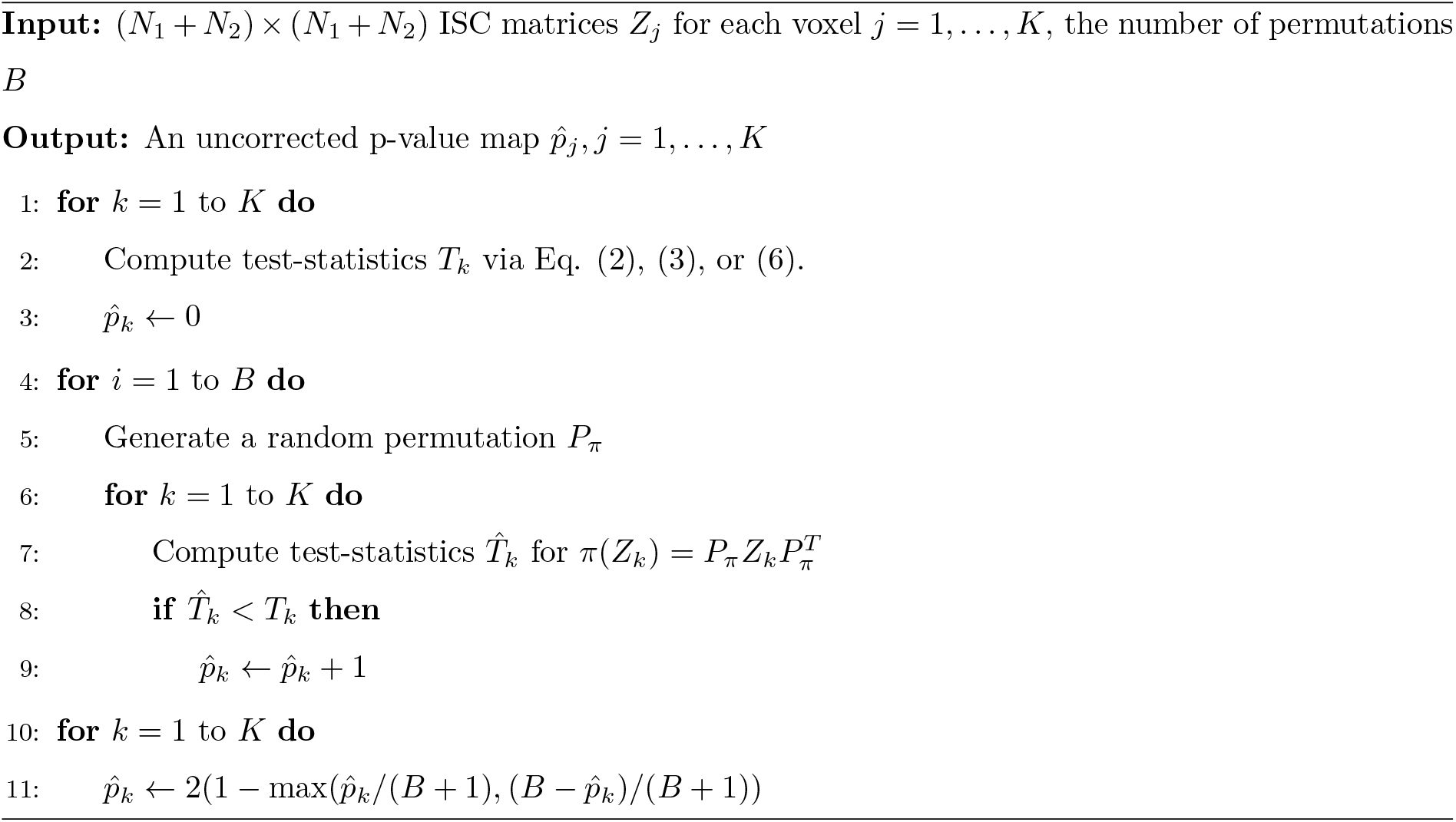

## Supporting information

Supplement: the ICBM-FRB subject information

## Appendix A Algorithms

## Appendix B False positive rates for null experiment by general linear model

Since the null-experiment (section 3.1) was based on the block-design data, we performed a group-difference test based on the standard General linear model (GLM) to help to verify that there were no significant group differences. This analysis was carried out using FSL’s (FMRIB’s Software Library, www.fmrib.ox.ac.uk/fsl) FEAT (FMRI Expert Analysis Tool) Version 6.00 using higher-level analysis of FLAME (FMRIB’s Local Analysis of Mixed Effects) stages 1+2 (Beckmann et al., 2003; Woolrich et al., 2004; Woolrich, 2008). The results of the analysis are in Table 6. These results verified that the standard GLM-based analysis produced false positive rates close to the nominal *α*-level, re-assuring that there was no group difference in fMRIs between the two matched groups of subjects.

**Table 6:** The results of the GLM-based analysis of the null-experiment of section 3.1. The columns *p* < 0.05, *p* < 0.01, and *p* < 0.001 list the fraction of the significant voxels at each *α* level (false positive rate). These results verify that also the standard GLM-based analysis produced false positive rates close to the nominal *α*-level, re-assuring that there was no group difference in fMRIs between the two matched groups of subjects.

| task | $p < 0.05$ | $p < 0.01$ | $p < 0.001$ |
| --- | --- | --- | --- |
| EO | 0.0469 | 0.0140 | 0.0030 |
| AN | 0.0474 | 0.0174 | 0.0036 |
| HA | 0.0172 | 0.0029 | 0.0004 |
| OM | 0.0700 | 0.0285 | 0.0063 |
| VG | 0.0402 | 0.0125 | 0.0023 |

### Algorithm 2 Global-null test with SW permutations (uncorrected)

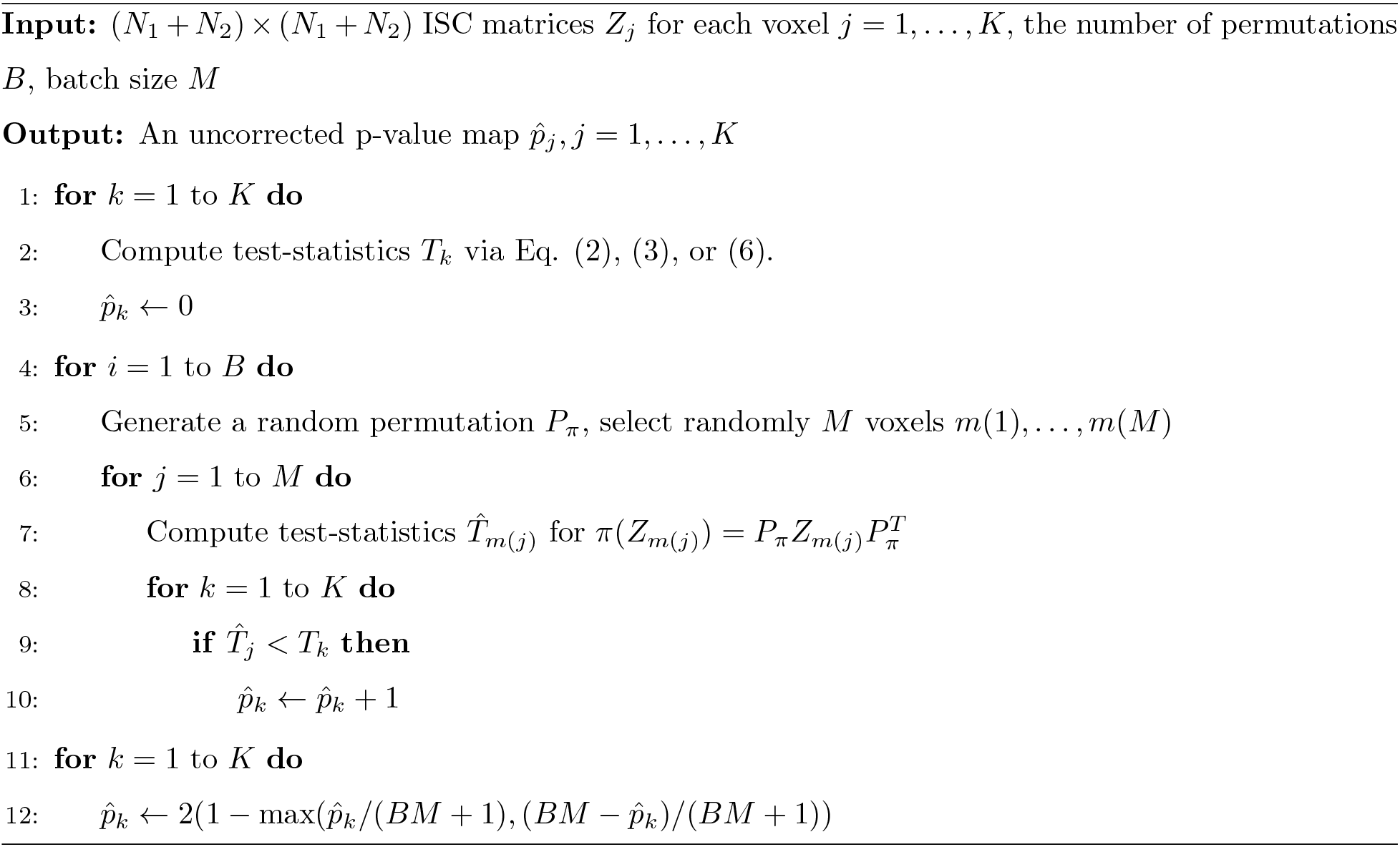

### Algorithm 3 Global-null test with SW permutations (FWE corrected)

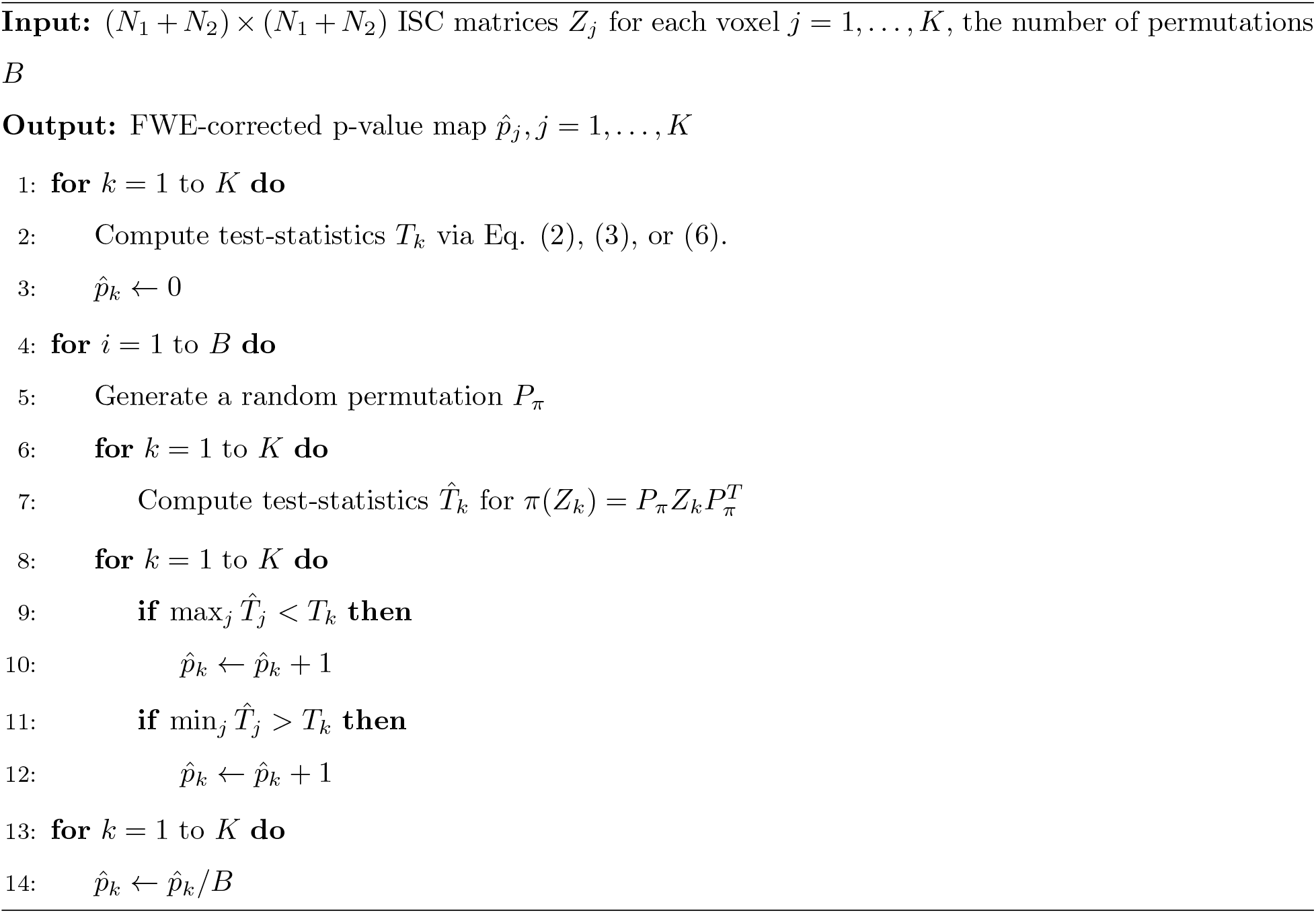

## Acknowledgments

J.T’s work is supported by the Academy of Finland (grant number 316258). J-P.K was funded by the Academy of Finland Postdoctoral Researcher program (Research Council for Natural Sciences and Engineering; grant number 286019). Data collection and sharing for this project was, in part, provided by the International Consortium for Brain Mapping (ICBM; Principal Investigator: John Mazziotta, MD, PhD). ICBM funding was provided by the National Institute of Biomedical Imaging and BioEngineering. ICBM data are disseminated by the Laboratory of Neuro Imaging at the University of Southern California. We thank Naree Kim and Seonhee Jang for use of the dance video and Paula Regener for use of the fMRI data comparing typical and autism spectrum observers.

## Footnotes

1 The correlation matrix of *X, corr(X)*, is a matrix of pairwise correlation coefficients between the columns of *X*. 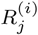 denotes the correlation matrix of 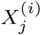. We apply Fisher’s z-transform to the elements of this matrix to obtain the z-transformed matrix: 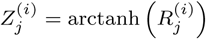.

