## Supplement: the ICBM-FRB subject information for "Comparing fMRI inter-subject correlations between groups using permutation tests"

| FRB-Orig | number | Anatomical | AuditoryNaming | VerbGeneration | ExternalOrder |
|---|---|---|---|---|---|
| Subject | 17 | ICBM_UCLA_1005 | ICBM_UCLA_10 | ICBM_UCLA_10 | ICBM_UCLA_1005 |
| Subject | 2 | ICBM_UCLA_1039 | ICBM_UCLA_10 | ICBM_UCLA_10 | ICBM_UCLA_1039 |
| Subject | 1 | ICBM_UCLA_1044 | ICBM_UCLA_10 | ICBM_UCLA_10 | ICBM_UCLA_1044 |
| Subject | 37 | ICBM_UCLA_1074 | ICBM_UCLA_10 | ICBM_UCLA_10 | ICBM_UCLA_1074 |
| Subject | 14 | ICBM_UCLA_1085 | ICBM_UCLA_10 | ICBM_UCLA_10 | ICBM_UCLA_1085 |
| Subject | 20 | ICBM_UCLA_1100 | ICBM_UCLA_11 | ICBM_UCLA_11 | ICBM_UCLA_1100 |
| Subject | 31 | ICBM_UCLA_1103 | ICBM_UCLA_11 | ICBM_UCLA_11 | ICBM_UCLA_1103 |
| Subject | 4 | ICBM_UCLA_1115 | ICBM_UCLA_11 | ICBM_UCLA_11 | ICBM_UCLA_1115 |
| Subject | 35 | ICBM_UCLA_1117 | ICBM_UCLA_11 | ICBM_UCLA_11 | ICBM_UCLA_1117 |
| Subject | 29 | ICBM_UCLA_1131 | ICBM_UCLA_11 | ICBM_UCLA_11 | ICBM_UCLA_1131 |
| Subject | 30 | ICBM_UCLA_1146 | ICBM_UCLA_11 | ICBM_UCLA_11 | ICBM_UCLA_1146 |
| Subject | 16 | ICBM_UCLA_1147 | ICBM_UCLA_11 | ICBM_UCLA_11 | ICBM_UCLA_1147 |
| Subject | 24 | ICBM_UCLA_1151 | ICBM_UCLA_11 | ICBM_UCLA_11 | ICBM_UCLA_1151 |
| Subject | 32 | ICBM_UCLA_1189 | ICBM_UCLA_11 | ICBM_UCLA_11 | ICBM_UCLA_1189 |
| Subject | 21 | ICBM_UCLA_1193 | ICBM_UCLA_11 | ICBM_UCLA_11 | ICBM_UCLA_1193 |
| Subject | 10 | ICBM_UCLA_1202 | ICBM_UCLA_12 | ICBM_UCLA_12 | ICBM_UCLA_1202 |
| Subject | 22 | ICBM_UCLA_1228 | ICBM_UCLA_12 | ICBM_UCLA_12 | ICBM_UCLA_1228 |
| Subject | 13 | ICBM_UCLA_1265 | ICBM_UCLA_12 | ICBM_UCLA_12 | ICBM_UCLA_1265 |
| Subject | 38 | ICBM_UCLA_1294 | ICBM_UCLA_12 | ICBM_UCLA_12 | ICBM_UCLA_1294 |
| Subject | 42 | ICBM_UCLA_1296 | ICBM_UCLA_12 | ICBM_UCLA_12 | ICBM_UCLA_1296 |
| Subject | 25 | ICBM_UCLA_1297 | ICBM_UCLA_12 | ICBM_UCLA_12 | ICBM_UCLA_1297 |
| Subject | 9 | ICBM_UCLA_1337 | ICBM_UCLA_13 | ICBM_UCLA_13 | ICBM_UCLA_1337 |
| Subject | 27 | ICBM_UCLA_1342 | ICBM_UCLA_13 | ICBM_UCLA_13 | ICBM_UCLA_1342 |
| Subject | 3 | ICBM_UCLA_1366 | ICBM_UCLA_13 | ICBM_UCLA_13 | ICBM_UCLA_1366 |
| Subject | 8 | ICBM_UCLA_1404 | ICBM_UCLA_14 | ICBM_UCLA_14 | ICBM_UCLA_1404 |
| Subject | 28 | ICBM_UCLA_1406 | ICBM_UCLA_14 | ICBM_UCLA_14 | ICBM_UCLA_1406 |
| Subject | 12 | ICBM_UCLA_1413 | ICBM_UCLA_14 | ICBM_UCLA_14 | ICBM_UCLA_1413 |
| Subject | 5 | ICBM_UCLA_1422 | ICBM_UCLA_14 | ICBM_UCLA_14 | ICBM_UCLA_1422 |
| Subject | 23 | ICBM_UCLA_1429 | ICBM_UCLA_14 | ICBM_UCLA_14 | ICBM_UCLA_1429 |
| Subject | 34 | ICBM_UCLA_1450 | ICBM_UCLA_14 | ICBM_UCLA_14 | ICBM_UCLA_1450 |
| Subject | 7 | ICBM_UCLA_1473 | ICBM_UCLA_14 | ICBM_UCLA_14 | ICBM_UCLA_1473 |
| Subject | 40 | ICBM_UCLA_1476 | ICBM_UCLA_14 | ICBM_UCLA_14 | ICBM_UCLA_1476 |
| Subject | 26 | ICBM_UCLA_1491 | ICBM_UCLA_14 | ICBM_UCLA_14 | ICBM_UCLA_1491 |
| Subject | 39 | ICBM_UCLA_1498 | ICBM_UCLA_14 | ICBM_UCLA_14 | ICBM_UCLA_1498 |
| Subject | 11 | ICBM_UCLA_1499 | ICBM_UCLA_14 | ICBM_UCLA_14 | ICBM_UCLA_1499 |
| Subject | 41 | ICBM_UCLA_1506 | ICBM_UCLA_15 | ICBM_UCLA_15 | ICBM_UCLA_1506 |
| Subject | 18 | ICBM_UCLA_1508 | ICBM_UCLA_15 | ICBM_UCLA_15 | ICBM_UCLA_1508 |
| Subject | 36 | ICBM_UCLA_1562 | ICBM_UCLA_15 | ICBM_UCLA_15 | ICBM_UCLA_1562 |
| Subject | 6 | ICBM_UCLA_1663 | ICBM_UCLA_16 | ICBM_UCLA_16 | ICBM_UCLA_1663 |
| Subject | 15 | ICBM_UCLA_1773 | ICBM_UCLA_17 | ICBM_UCLA_17 | ICBM_UCLA_1773 |
| Subject | 19 | ICBM_UCLA_1795 | ICBM_UCLA_17 | ICBM_UCLA_17 | ICBM_UCLA_1795 |
| Subject | 33 | ICBM_UCLA_1907 | ICBM_UCLA_19 | ICBM_UCLA_19 | ICBM_UCLA_1907 |

removed

1246
1217
1158
1155

ICBM_UCLA_subject_numbers_vs_data

| HandImitation | Oculomotor | | | M | F | age | hand | | |
|---|---|---|---|---|---|---|---|---|---|
| | | | | | | | R | L | |
| ICBM_UCLA_1005 | ICBM_UCLA_1005 | Subject | 17 | 1 | | 23 | 1 | | |
| ICBM_UCLA_1039 | ICBM_UCLA_1039 | Subject | 2 | 1 | | 23 | 1 | | |
| ICBM_UCLA_1044 | ICBM_UCLA_1044 | Subject | 1 | 1 | | 31 | 0 | | A |
| ICBM_UCLA_1074 | ICBM_UCLA_1074 | Subject | 37 | 1 | | 33 | 1 | | |
| ICBM_UCLA_1085 | ICBM_UCLA_1085 | Subject | 14 | 1 | | 33 | 1 | | |
| ICBM_UCLA_1100 | ICBM_UCLA_1100 | Subject | 20 | 1 | | 23 | 1 | | |
| ICBM_UCLA_1103 | ICBM_UCLA_1103 | Subject | 31 | 1 | | 34 | 1 | | |
| ICBM_UCLA_1115 | ICBM_UCLA_1115 | Subject | 4 | 1 | | 31 | 1 | | |
| ICBM_UCLA_1117 | ICBM_UCLA_1117 | Subject | 35 | 1 | | 32 | 1 | | |
| ICBM_UCLA_1131 | ICBM_UCLA_1131 | Subject | 29 | 1 | | 21 | 1 | | |
| ICBM_UCLA_1146 | ICBM_UCLA_1146 | Subject | 30 | 0 | F | 33 | 1 | | |
| ICBM_UCLA_1147 | ICBM_UCLA_1147 | Subject | 16 | 0 | F | 32 | 1 | | |
| ICBM_UCLA_1151 | ICBM_UCLA_1151 | Subject | 24 | 1 | | 22 | 1 | | |
| ICBM_UCLA_1189 | ICBM_UCLA_1189 | Subject | 32 | 1 | | 26 | 0 | L | |
| ICBM_UCLA_1193 | ICBM_UCLA_1193 | Subject | 21 | 0 | F | 23 | 1 | | |
| ICBM_UCLA_1202 | ICBM_UCLA_1202 | Subject | 10 | 0 | F | 21 | 1 | | |
| ICBM_UCLA_1228 | ICBM_UCLA_1228 | Subject | 22 | 1 | | 22 | 0 | L | |
| ICBM_UCLA_1265 | ICBM_UCLA_1265 | Subject | 13 | 0 | F | 37 | 0 | L | |
| ICBM_UCLA_1294 | ICBM_UCLA_1294 | Subject | 38 | 0 | F | 30 | 1 | | |
| ICBM_UCLA_1296 | ICBM_UCLA_1296 | Subject | 42 | 0 | F | 30 | 1 | | |
| ICBM_UCLA_1297 | ICBM_UCLA_1297 | Subject | 25 | 0 | F | 25 | 1 | | |
| ICBM_UCLA_1337 | ICBM_UCLA_1337 | Subject | 9 | 0 | F | 22 | 1 | | |
| ICBM_UCLA_1342 | ICBM_UCLA_1342 | Subject | 27 | 1 | | 31 | 1 | | |
| ICBM_UCLA_1366 | ICBM_UCLA_1366 | Subject | 3 | 0 | F | 28 | 1 | | |
| ICBM_UCLA_1404 | ICBM_UCLA_1404 | Subject | 8 | 1 | | 28 | 0 | L | |
| ICBM_UCLA_1406 | ICBM_UCLA_1406 | Subject | 28 | 0 | F | 32 | 1 | | |
| ICBM_UCLA_1413 | ICBM_UCLA_1413 | Subject | 12 | 1 | | 22 | 1 | | |
| ICBM_UCLA_1422 | ICBM_UCLA_1422 | Subject | 5 | 1 | | 36 | 1 | | |
| ICBM_UCLA_1429 | ICBM_UCLA_1429 | Subject | 23 | 1 | | 21 | 1 | | |
| ICBM_UCLA_1450 | ICBM_UCLA_1450 | Subject | 34 | 1 | | 25 | 1 | | |
| ICBM_UCLA_1473 | ICBM_UCLA_1473 | Subject | 7 | 1 | | 21 | 1 | | |
| ICBM_UCLA_1476 | ICBM_UCLA_1476 | Subject | 40 | 1 | | 32 | 1 | | |
| ICBM_UCLA_1491 | ICBM_UCLA_1491 | Subject | 26 | 0 | F | 23 | 1 | | |
| ICBM_UCLA_1498 | ICBM_UCLA_1498 | Subject | 39 | 0 | F | 32 | 1 | | |
| ICBM_UCLA_1499 | ICBM_UCLA_1499 | Subject | 11 | 0 | F | 25 | 1 | | |
| ICBM_UCLA_1506 | ICBM_UCLA_1506 | Subject | 41 | 0 | F | 36 | 1 | | |
| ICBM_UCLA_1508 | ICBM_UCLA_1508 | Subject | 18 | 0 | F | 27 | 1 | | |
| ICBM_UCLA_1562 | ICBM_UCLA_1562 | Subject | 36 | 0 | F | 35 | 1 | | |
| ICBM_UCLA_1663 | ICBM_UCLA_1663 | Subject | 6 | 1 | | 36 | 1 | | |
| ICBM_UCLA_1773 | ICBM_UCLA_1773 | Subject | 15 | 1 | | 32 | 1 | | |
| ICBM_UCLA_1795 | ICBM_UCLA_1795 | Subject | 19 | 0 | F | 33 | 1 | | |
| ICBM_UCLA_1907 | ICBM_UCLA_1907 | Subject | 33 | 0 | F | 27 | 1 | | |
| | | | | | awg | 28.3095238095238 | | | |
| | | | | | M: | 23 | R: | 37 | |
| | | | | | F: | 19 | L: | 4 | |

| Group |
|---|
| 2 |
| 2 |
| 2 |
| 1 |
| 1 |
| 1 |
| 2 |
| 1 |
| 1 |
| 2 |
| 1 |
| 1 |
| 2 |
| 1 |
| 2 |
| 2 |
| 1 |
| 2 |
| 2 |
| 1 |
| 1 |
| 1 |
| 2 |
| 2 |
| 1 |
| 2 |
| 1 |
| 1 |
| 2 |
| 2 |
| 2 |
| 2 |
| 1 |
| 2 |
| 1 |
| 1 |
